# Early delta cortical tracking to non-verbal auditory stimuli predicts reading skills: A one-year longitudinal study

**DOI:** 10.1101/2023.10.04.560887

**Authors:** Camila Zugarramurdi, Lucía Fernández, Marie Lallier, Juan Carlos Valle-Lisboa, Manuel Carreiras

## Abstract

The precision of cortical tracking of auditory rhythmic stimuli in low frequency ranges coding for prosodic (delta) and syllabic (theta) amplitude modulations in the speech signal has been proposed to contribute to the development of phonological processing and reading acquisition. The present study investigates the role of low frequency cortical tracking of non-verbal auditory stimuli in reading acquisition through a longitudinal design from before the onset of formal reading instruction until one year afterwards. At time one, 40 prereading children performed a passive listening task of amplitude modulated white noise presented in the delta (2 Hz) and theta (4 Hz) frequency bands, while their neural activity was recorded via EEG. At time two, at the end of first grade, children’s reading skills were assessed. Results show significant cortical tracking in prereading children at both frequencies, with larger responses for theta than for delta rate non-verbal auditory stimuli. Importantly, only pre-reding cortical tracking measures for delta rate stimuli predicted reading acquisition one year later. These findings underscore the role of early neural synchronization to delta rate rhythmic auditory stimuli to reading acquisition and support the potential important role of early prosodic processing for the development of future reading skills.

---

Reading skills play a major role in personal and professional development in modern societies (Huettig & Pickering, 2019). The acquisition of reading skills, especially during its early stages, strongly relies on oral language abilities, including efficient and precise speech processing. At the auditory level, speech processing is achieved through cortical tracking (CT). Cortical tracking to speech refers to the modulation of cortical oscillations in response to acoustic onset edges in the speech envelope (Oganian & Chang, 2019). Cortical oscillations in the auditory cortex fluctuate at preferred frequencies at delta, theta and gamma rates (1-3 Hz, 4-7 and above 30 Hz, respectively), which broadly match the temporal fluctuations present in the speech signal, corresponding to prosodic, syllabic and phonemic linguistic information, respectively (Peelle & Davis, 2012). Regarding its functional role, CT of the speech envelope is necessary for speech comprehension; when the speech envelope is disrupted, CT is reduced and intelligibility is hindered (Ahissar et al., 2001). Since CT underlies speech processing, it could be expected that disruptions in CT would result in phonological deficits and, in turn, in reading difficulties (Goswami, 2011).

Several studies have tested the association between CT and reading skills by comparing CT to auditory stimuli between dyslexic and typical readers. In adults, studies have shown differences in CT between dyslexics and controls in the delta (Hämäläinen et al., 2012), theta (Lizarazu et al., 2015) and gamma (Lehongre et al., 2011; Lizarazu et al., 2015) bands, using linguistic and non-linguistic stimuli. In children, differences in CT have also been observed in delta (Cutini et al., 2016; Lallier et al., 2016; Power et al., 2013, 2016), theta (Lizarazu et al., 2015) and gamma (Lehongre et al., 2011). While the above evidence suggests CT to auditory stimuli might be related to reading skill, evidence is still inconclusive. Critically, a recent large study which aimed to replicate the previous findings mostly failed to do so (Lizarazu et al., 2021). Interpretation of the previous studies is further complicated by the fact that most employed samples that had already acquired reading skills. Therefore, any observed difference between groups might be due to the reduced reading experience of dyslexics rather than a cause of reading difficulties, given it is by now well established that reading experience modifies speech processing (Araújo et al., 2018; Castles & Coltheart, 2004; Huettig et al., 2018).

In the present study we provide novel evidence through a longitudinal approach regarding the role of CT in reading acquisition. We examined CT to non-linguistic stimuli in prereading children at midterm kindergarten, and their reading development one year later, after they had received 7 to 9 months of reading instruction. Our hypothesis was that CT at the delta and/or theta rate measured in prereaders would correlate to reading acquisition one year later, in line with the temporal sampling framework (Goswami, 2011) and recent empirical evidence (De Vos et al., 2017; Ríos-López et al., 2020; Woodruff Carr et al., 2014). To test this hypothesis, through a passive listening paradigm, we computed CT at speech relevant rates (delta and theta) and related it to reading skills one year later. By using non-linguistic stimuli, we aimed to test whether the mechanisms and relevant frequencies of cortical tracking and its relation to reading are dependent on the verbal nature of the stimuli, and to control for variations in the broadband energy of the stimuli at different frequencies.

## Methods

### Sample

Forty children attending kindergarten (K5) took part in the study (21 males, age range 5 – 6.5 years, mean = 6.1). These were selected as part of a larger study on reading acquisition (reported in Zugarramurdi et al., 2021, 2022). All participants’ caregivers provided informed written consent and all children verbally agreed to participate. All participants were white, native Spanish speakers, had normal or corrected-to-normal vision and reported no hearing impairments. Behavioural data were collected between June and August during kindergarten year (mid-term, since the school year starts in March), and between October and December while in first grade (end of term). Electrophysiological data were collected between November and February during kindergarten year (end of term). Data from four children were discarded due to excess noise (two children) and technical issues during recording (two children), meaning the final sample was composed of 36 children.

### Behavioural measures

During kindergarten, reading status was assessed by asking children to read aloud a list of 15 high frequency words and 15 pseudowords printed on paper. Since no standardized norms were available for this population, a list of high frequency words was created (see Supplementary Material for the complete list). During first grade, reading skills were assessed by asking children to read aloud a list of 30 words and 30 pseudowords, in a tablet, one word per screen (see Supplementary Material for the complete list). Nonverbal IQ was measured using the Matrix Reasoning subtest of the Spanish version of the Wechsler Preschool and Primary Scale of Intelligence (Wechsler, 2001).

### Neural measures

#### Stimuli

Stimuli consisted of amplitude modulated white noise in 3 conditions: 2, 4 and 8 Hz with 100% depth, and a non-modulated condition (Lizarazu et al., 2015). The 8 Hz condition was used as a control for a non speech-relevant rate, and the unmodulated condition was used as a control for the modulation manipulation itself. Each condition was presented 24 times in 10 second trials. Trials were presented in random order with no inter-trial-interval. The whole stimuli lasted 16 minutes. Stimuli presentation was coded in PsychoPy (Peirce, 2008), using the sound library with *pyo* backend in Windows 7.

Stimuli were presented binaurally through Etymotic ETY Kids 5 insert earphones. Sound pressure level adjustment for each child consisted of listening to a recorded sentence (*Donde viven los monstruos* [Where monsters live]) and repeating it correctly. Throughout the session, children viewed silent cartoons projected onto the wall in order to keep them entertained and as quiet as possible. The session was interrupted if children showed signs of boredom or tiredness.

#### EEG recording and processing

##### Recording

EEG data were acquired using a Biosemi Active Two system, with 32 electrodes in a 20-10 layout. Activity was referenced online to the common mode sense (CMS, active electrode) and grounded to a passive electrode (Driven Right Leg, DRL). Data were digitized at 512 Hz.

##### Pre-processing

The EEG signal was processed using Fieldtrip toolbox (Oostenveld et al., 2011) in Matlab R2018a (The Mathworks Inc., 2018) and custom code. In the pre-processing step, the continuous EEG signal was band passed with a two-pass fourth-order Butterworth filter between 0.1 and 40 Hz, baselined to 0.8 s prior to stimulus onset and re-referenced to the Cz electrode. In each 10 second trial, the first second was discarded due to an observed increase in noise in response to the incoming stimuli, and the signal was redefined into 2 second epochs with 1 second overlap for all conditions. Within each 2 second epoch, artifact rejection was based on an adaptation of Junghöfer et al (2000). In each stimulus condition, individual channels and epochs were rejected if they surpassed a threshold defined by the median of the standard deviation for all channels/trials according to the following equation (Fló et al., 2019):

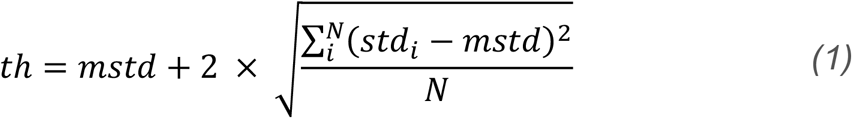

Rejected channels were interpolated using the default “weighted” method in the ft_channelrepair function and a Biosemi 32 template for defining neighbours with the ft_prepare_neighbours function. According to the template used for neighbour selection, each channel has on average 6 neighbours, ranging from 3 to 8 according to channel position. On average, after artifact rejection, there were 140 epochs per child per condition.

Next, power estimates were obtained through the discrete Fourier transform. For each condition, 2-second-epochs were concatenated in groups of 5 in order to increase spectral resolution. The number of epochs was standardized across children by limiting it to a range between 90 and 120; for children with more than 120 epochs, subsequent epochs were discarded. Thus, for each child there were between 18 and 24 sweeps per condition. Next, sweeps were averaged in the time domain and transformed into the frequency domain. Signal was padded with zeros to the next power of two in order to improve performance.

##### Signal-to-noise ratio

The previous processing steps yielded power estimates per child per channel per condition with a spectral resolution of 0.0625 Hz (Figure *1*.A). Signal-to-noise ratio (SNR) was used to quantify the degree of synchronized neural activity (De Vos et al., 2020). SNR was computed for each stimulus frequency as:

**Figure 1.**
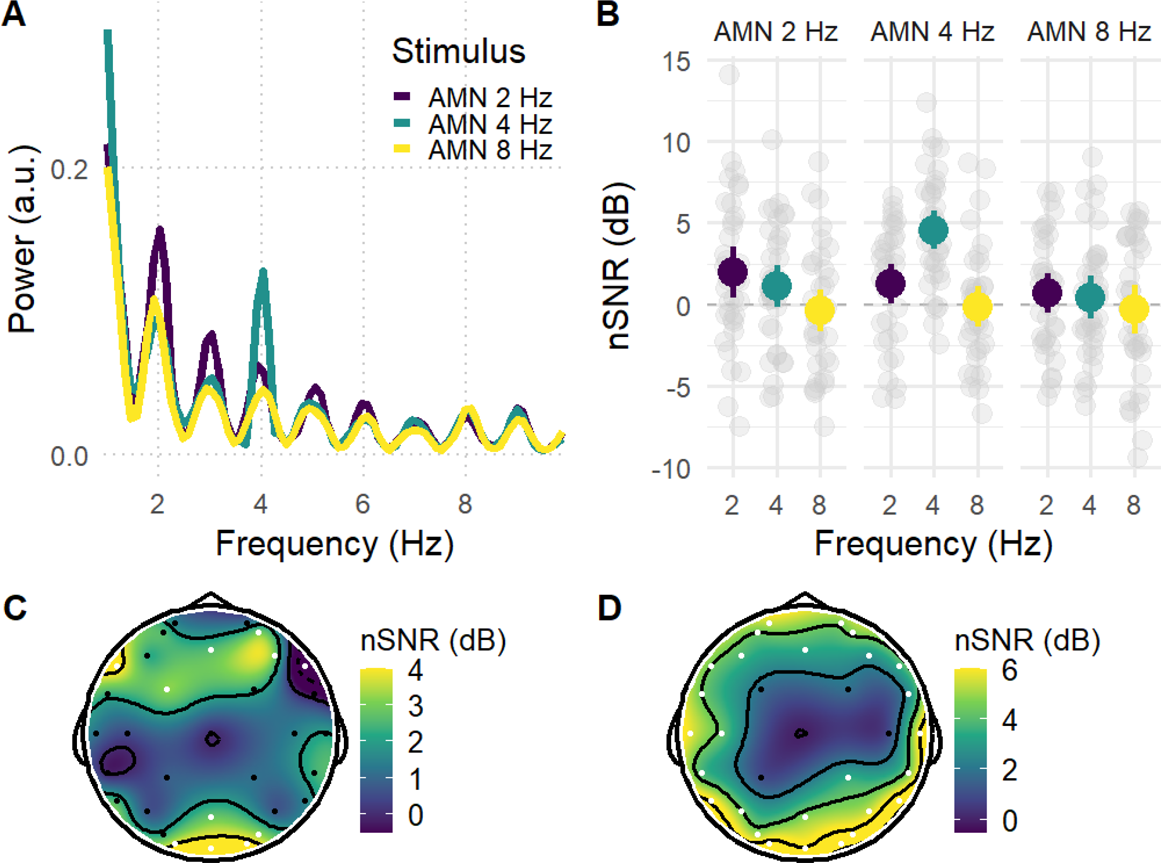
A. Power spectrum of the EEG response for each stimulus condition. Peaks are observed when the stimulus frequency matches the response frequency at 2 and 4 Hz. B. SNR responses averaged across electrodes for all stimuli and response frequencies. Dots represent each child’s individual average across electrodes; C. Topographical distribution of CT at delta rate. Two clusters of CT can be observed, one anterior and one posterior. D. Topographical distribution of CT at theta rate. A scalp-wide CT is observed. AMN: amplitude modulated noise. nSNR: normalized signal to noise ratio. CT: cortical tracking.

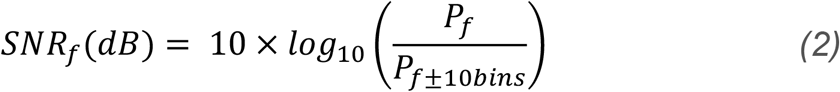

where *P*_*f*_is the response power, and *P*_*f*±10*bins*_is the power in the 10 adjacent bins to the stimulus frequency.

Next, in order to obtain the specific response to the stimulus of interest, normalized SNR was defined as the subtraction of the SNR for each stimulus frequency from the SNR for unmodulated stimuli (control condition), as follows:

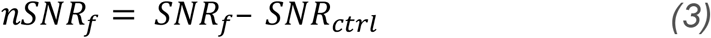

For each AMN frequency, 3 normalized SNR were computed corresponding to the 3 stimulus frequencies. This resulted in a 3 x 3 design of AMN stimulus frequency (2, 4 and 8 Hz) x nSNR frequency (2, 4 and 8 Hz).

CT was defined as instances when the response to a stimulus of interest was significantly larger that the response observed for the unmodulated control condition, i.e., a nSNR significantly larger than zero. Upper and lower thresholds were defined for each stimulus condition as *M±3*SD*, trials with nSNRs values outside of this threshold were replaced by the threshold value (0.8% of the trials).

### Statistical analysis

To predict brain responses, linear mixed effects models were built using the *lme4* package in R (Bates et al., 2015; R Core Team, 2018). For all models, planned comparisons were conducted through the *emmeans* package. Degrees of freedom were estimated through the Satterthwaite method, and p-values were adjusted for multiple comparisons through the false discovery rate (FDR) method.

## Results

First, CT to the stimulus conditions for each response frequency was examined. Next, the topographical distribution of the response was investigated. In order to avoid ad-hoc electrode selection criteria, CT was examined in an overarching model with all electrodes. Finally, the relation between CT and reading acquisition was investigated.

### CT to AMN

A linear mixed effects model was computed with nSNR as outcome and main effects of stimulus frequency and response frequency and their interaction as predictors. Random intercepts for subject were included in order to account for the repeated measures of electrodes over subjects. Model results showed main effects of stimulus frequency and response frequency as well as a significant interaction (all p < 0.001). Estimated marginal means and planned comparisons were obtained for each stimulus and all response frequencies for all main effects and the interaction (Table 1). An estimated marginal mean for nSNR larger than zero was interpreted as a specific CT to the target stimuli (i.e., significantly larger CT to AMN than to unmodulated noise stimuli).

**Table 1.**
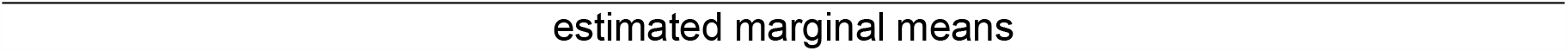
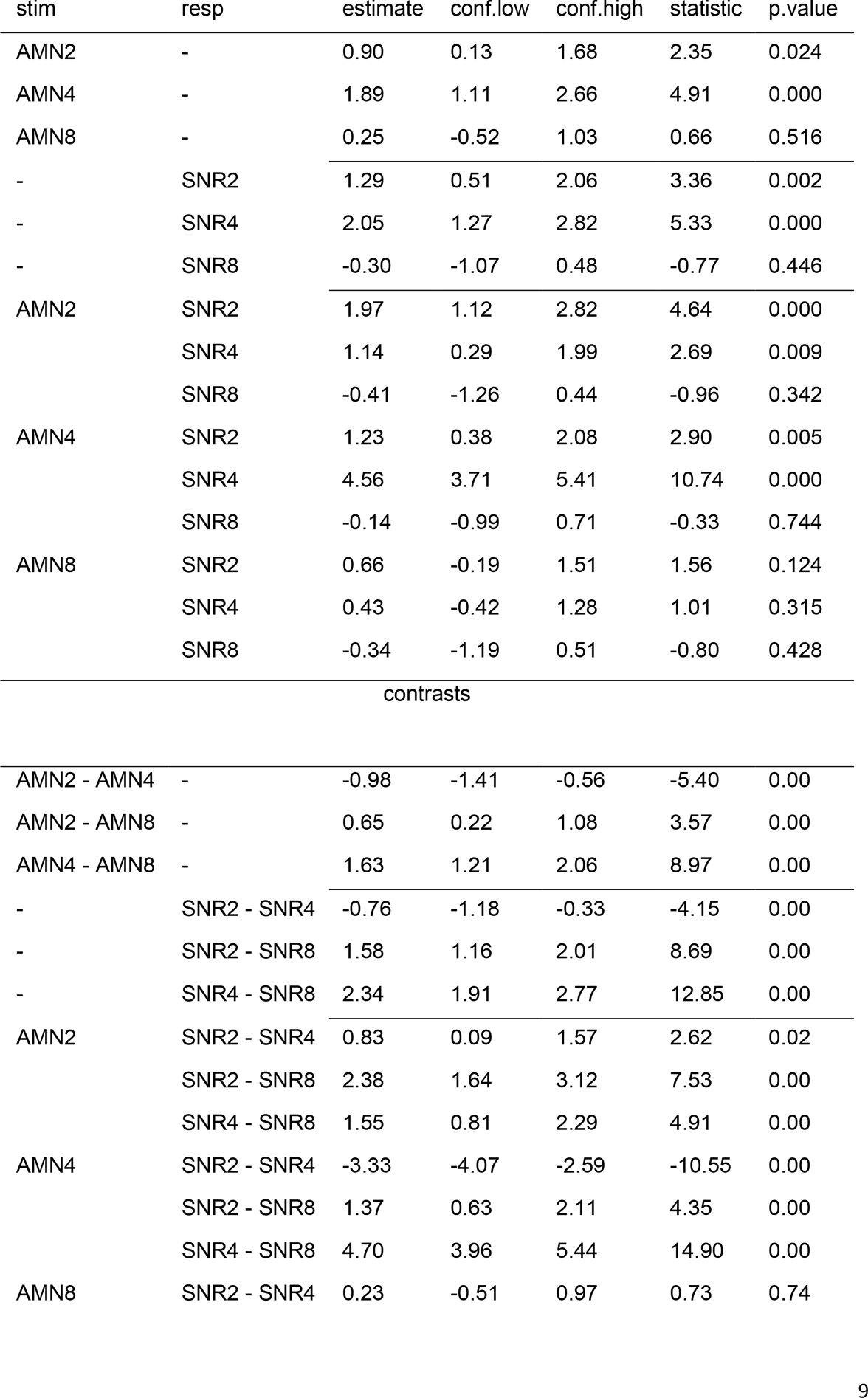
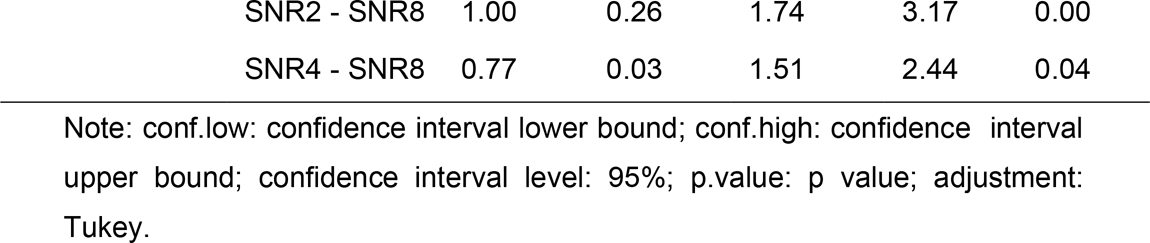
Estimated marginal means and their contrasts for each stimulus and all response frequencies.

With respect to *stimulus* frequencies, CT was found for AMN stimuli at the speech relevant rates of 2 and 4 Hz but not for the control condition of 8 Hz (Figure 1. B). CT differed across all pairwise comparisons, with responses to the 4Hz stimuli being largest, followed by responses to the 2 Hz and 8 Hz stimuli (Table 1). The same pattern was observed with respect to *response* frequencies: CT was found for 2 and 4 Hz responses, but not for 8 Hz, and differed across all pairwise comparisons, with 4Hz responses being largest, followed by 2 Hz and 8 Hz.

When looking at the interaction between *stimulus and response* frequencies, results showed that, for each stimulus frequency, responses were largest when the stimulus frequency matched the response frequency (Table 1). In other words, when children heard 4 Hz stimuli, they showed larger responses at 4 Hz than at 2 or 8 Hz; and when they heard 2 Hz stimuli, they showed larger responses at 2 Hz than at 4 or 8 Hz. This was not the case in the 8 Hz stimuli, where none of the responses to AMN were significantly different from the unmodulated condition.

Thus, results showed that brain responses tune to auditory stimuli at frequencies relevant to the speech input (2 and 4 Hz). Furthermore, CT was largest for response frequencies corresponding to the stimulus frequency. In order to investigate the topographical distribution of CT, we further explored CT by electrode.

### Topography

The topographical distribution of CT was studied for delta and theta stimuli (2 and 4 Hz, respectively; Figure 1.C.). Two linear mixed effects models were fit (one for each frequency) with CT as outcome and electrode as predictor. Random intercepts for children were included to account for the repeated measures of electrodes. Both frequencies showed a significant effect of electrode (delta: *X*^*2*^(30) = 49.4, p = 0.014; theta: *X*^*2*^(30) = 81.7, p < 0.001), so t-tests for each electrode were computed, with false-discovery rate correction for multiple comparisons. For delta, two clusters of CT were observed, one anterior (AF4, F7, F4, FC1, Fz) and one posterior (PO4, O1, O2, Pz, Oz). A significant F8 response was also observed, but SNR was negative, meaning the response to the unmodulated stimuli was larger than to the modulated stimuli. It is unclear whether this incongruent effect was due to artifact. After false discovery rate correction was applied for multiple comparisons, CT remained significant at one anterior site (F7) and at three posterior sites (O1, O2, and Oz). In contrast, for theta, a broad scalp-wide CT was observed, except for four central electrodes (FC1, FC2, C4 and CP1, FDR corrected).

### CT and future reading skills

Next, we tested whether CT at each frequency was related to reading acquisition one year later. During kindergarten, most children (n = 33) could not read any of the presented words and pseudowords, and three children had already started learning how to read (they could read 9 out of 15 pseudowords, on average). At the end of first grade, mean reading accuracy was 0.75 (SD = 0.32, min = 0, max = 1) and reading scores displayed a bimodal distribution (see Zugarramurdi et al., 2021 for a broader description). Based on a -1 z score threshold, children were divided into two groups: *poor readers* (PR, mean accuracy = 0.13, min = 0, max = 0.45, n = 7) and *typical readers* (TR, mean accuracy = 0.89, min = 0.7, max = 1, n = 29).

We computed two linear mixed effects regression models —one for delta and one for theta— with CT during kindergarten as outcome, and fixed effects for reading group at the end of first grade (PR vs. TR), electrode (delta: n = 4; theta: n = 27) and their interaction as predictors. Random intercepts for children were included to account for the repeated measures of electrodes within children. Levene’s test was used to check for homogeneity of variance, which was met for both models (delta: F(1,142) = 0.002, p = 0.96); theta: F(1,970) = 1.03 p = 0.3 (Foster et al., 2011)).

For delta, a main effect of reading (X2(1) = 5.66, p = 0.017), a marginal main effect of electrode (X2(3) = 7.29, p = 0.063), and a significant interaction were observed (X2(3) = 8.79, p = 0.032). Post-hoc tests showed that poor readers did not show significant CT to AMN (mean nSNR = 2.95, SE = 2.2, 95% CI = [-2.22, 8.12], t = 1.33, pFDR = 0.18), whereas typical readers did (mean nSNR = 4.59, SE = 1.08, 95% CI = [2.06, 7.13], t = 4.24, pFDR = 0.0003) (Figure 2). Overall contrasts did not show differences between groups (PR – TR estimate = -1.64, SE = 2.46, 95% CI = [-6.63, 3.35], t = -0.67, pFDR = 0.50). However, when electrodes were included, significant differences between groups were observed at the F7 electrode but not in the occipital cluster (Table 2). A linear model predicting CT at F7 from reading group, Age, and IQ, showed that reading group predicted CT above and beyond Age and IQ (model F(3,32) = 2.83, p = 0.053; reading group coefficient: estimate = 7.89, SE = 3, t = 2.65, p= 0.013).

**Table 2.**
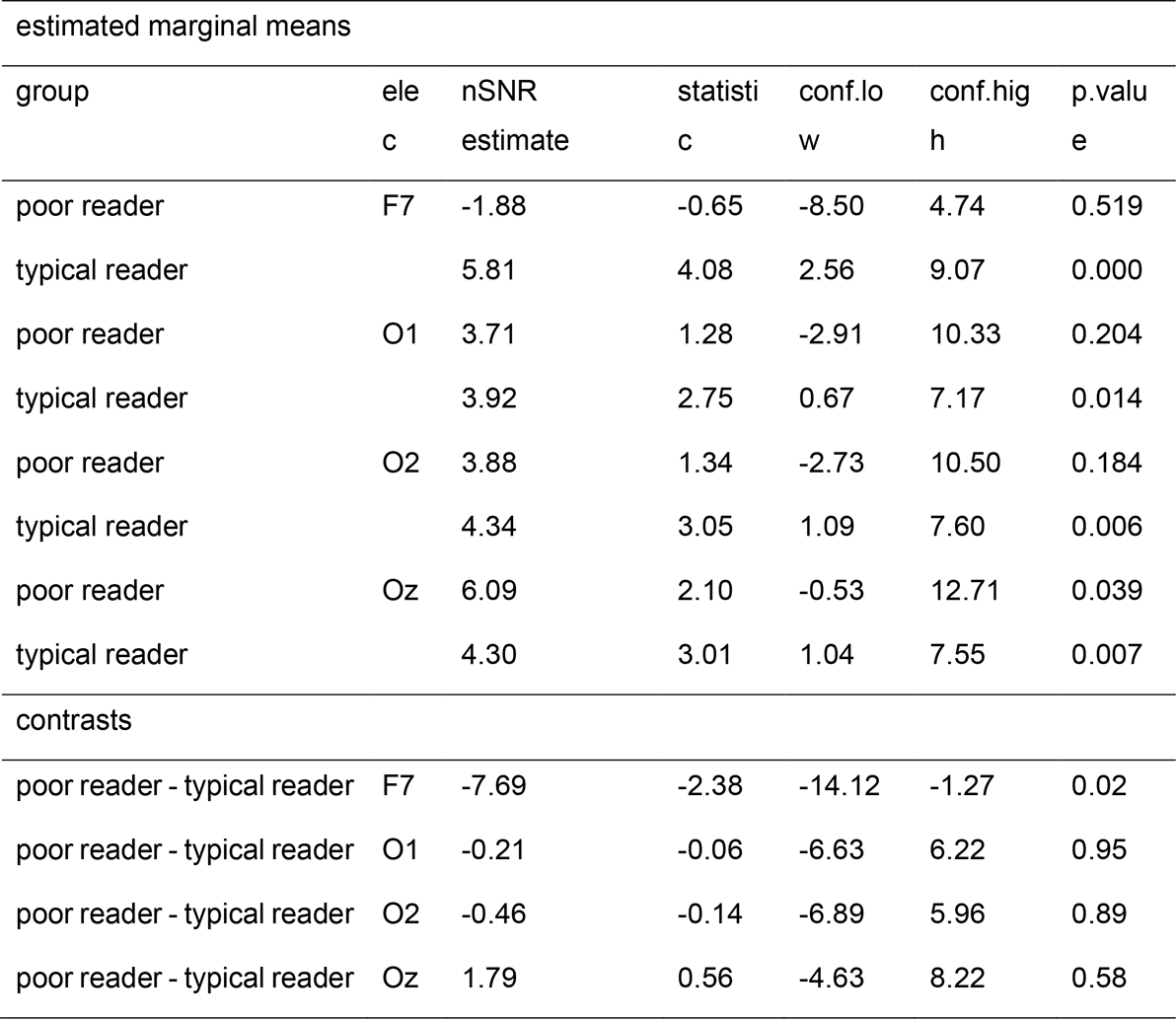
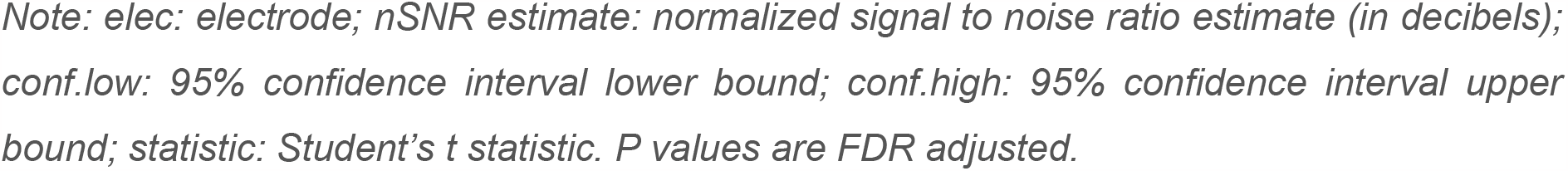
Delta neural synchronization at each electrode and frequency, and contrasts between reading groups.

| estimated marginal means |  |  |  |  |  |  |
| --- | --- | --- | --- | --- | --- | --- |
| group | elec | nSNR estimate | statistic | conf.lo w | conf.hig h | p.value |
| poor reader | F7 | -1.88 | -0.65 | -8.50 | 4.74 | 0.519 |
| typical reader |  | 5.81 | 4.08 | 2.56 | 9.07 | 0.000 |
| poor reader | O1 | 3.71 | 1.28 | -2.91 | 10.33 | 0.204 |
| typical reader |  | 3.92 | 2.75 | 0.67 | 7.17 | 0.014 |
| poor reader | O2 | 3.88 | 1.34 | -2.73 | 10.50 | 0.184 |
| typical reader |  | 4.34 | 3.05 | 1.09 | 7.60 | 0.006 |
| poor reader | Oz | 6.09 | 2.10 | -0.53 | 12.71 | 0.039 |
| typical reader |  | 4.30 | 3.01 | 1.04 | 7.55 | 0.007 |
| contrasts |  |  |  |  |  |  |
| poor reader - typical reader | F7 | -7.69 | -2.38 | -14.12 | -1.27 | 0.02 |
| poor reader - typical reader | O1 | -0.21 | -0.06 | -6.63 | 6.22 | 0.95 |
| poor reader - typical reader | O2 | -0.46 | -0.14 | -6.89 | 5.96 | 0.89 |
| poor reader - typical reader | Oz | 1.79 | 0.56 | -4.63 | 8.22 | 0.58 |

Note: *elec*: electrode; *nSNR estimate*: normalized signal to noise ratio estimate (in decibels); *conf.low*: 95% confidence interval lower bound; *conf.high*: 95% confidence interval upper bound; *statistic*: Student's *t* statistic. *P* values are FDR adjusted.

**Figure 2.**
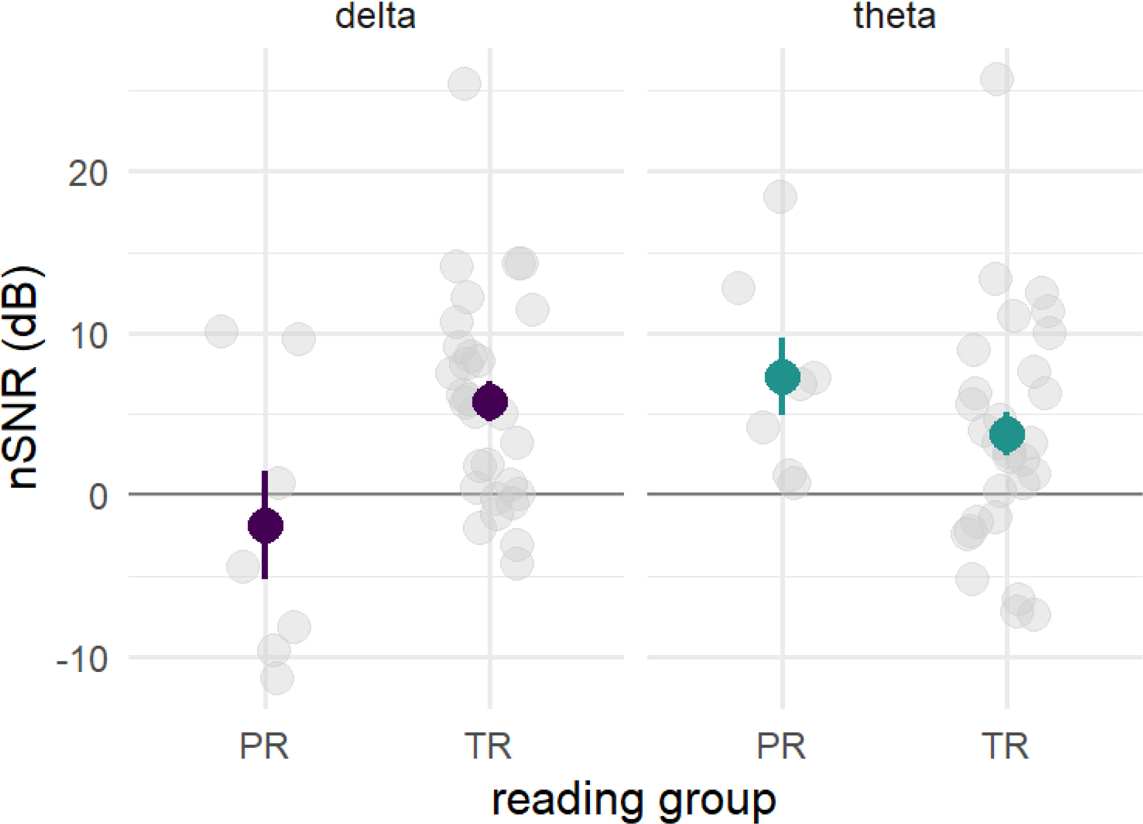
Neural synchronization by reading group at delta and theta rates at F7 electrode. Error bars indicate standard error of the mean. PR: poor reader; TR: typical reader. Dots represent individual children.

## Discussion

### CT in prereaders

Our findings show that prereading children show CT to auditory stimuli at both delta and theta rates (but not at 8 Hz), thus showing a cerebral specialization for auditory processing at speech-relevant rates. Although previous studies with prereaders have reported CT to theta rate (Vanvooren et al., 2014) and delta rate (Ríos-López et al., 2020, 2022), neither of these studies found CT to both rates, as shown here. While Vanvooren et al. did not analyse delta rate, Ríos-López et al. tested both frequencies in response to speech and failed to find CT at theta. Studies with infants have also shown CT at delta (Attaheri et al., 2020) and theta (Ortiz Barajas et al., 2021) in response to infant-directed speech. Thus, our findings contribute to the emerging picture suggesting CT to auditory stimuli at speech-relevant rates develops very early on, possibly in a continuous manner. By comparing CT to delta and theta frequencies in response to amplitude modulated white noise, we were able to show that CT to non-linguistic stimuli was stronger and more widely distributed for theta than delta rate. In contrast to speech, the non-linguistic stimulus used allowed us to control for differences in spectral energy between frequencies, suggesting that the differences in the response do not stem from differences in the spectral energy of the frequency band.

For theta CT, the large scalp-wide distribution was somewhat surprising. Previous studies have reported theta CT at temporo-parietal electrodes with fNIRS (Cutini et al., 2016), EEG (Lehongre et al., 2013; Vanvooren et al., 2014) and MEG (Palana et al., 2022), broadly corresponding to auditory cortex in the temporal lobe. However, source reconstruction of auditory steady state responses to 40 Hz AMN stimuli found sources in central auditory pathways —both cortical and subcortical, including brainstem and primary auditory cortex— and extra auditory pathways broadly distributed —including pre and post central gyri, orbitofrontal, parahippocampal, occipital, superior parietal and cingulate gyri (Farahani et al., 2020). When the stimulus was limited to the theta range, sources have been found in the frontal lobe and medial limbic structures (Farahani et al., 2017) and in associative auditory and non-auditory cortex (Giraud et al., 2000). Thus, it is possible that the observed topographical distribution results from our data-driven approach to electrode selection, and that it originates from both auditory and extra-auditory sources.

For delta CT, we found a frontal and an occipital cluster, as opposed to the expected temporo-parietal distribution (e.g. Lehongre et al., 2013). However, delta CT has also been described in both auditory and non-auditory cortices. On one hand, delta CT has been observed in the auditory cortex, involved in bottom-up segmentation of the speech input (Ghitza, 2017; Molinaro et al., 2016). On the other hand, delta CT has also been shown in the frontal lobe — in particular IFG and precentral gyrus— exerting top-down modulations on delta and theta activity of the auditory cortex (Park et al., 2015). These top-down modulations have been implicated in the grouping of words into syntactic phrases (Ding et al., 2015; Meyer et al., 2019), temporal predictions during speech processing (Rimmele et al., 2018), sensory chunking of articulated sounds (Heisz et al., 2009), and attentional modulation (Wild et al., 2012). The observed distribution in the current data is more compatible with top-down frontal effects than bottom-up temporal processing. Moreover, since our stimuli were non-linguistic, it is more likely to reflect non-linguistic mechanisms such as temporal prediction, sensory chunking, or attentional modulation. Importantly, we failed to find neural delta CT at the auditory level. At present, it is hard to tell whether this reflects developmental differences or if it is a consequence of our experimental paradigm or data processing.

### CT and future reading skills

With respect to CT and reading, results showed that differences in delta —but not theta— tracking, precede and predict future reading acquisition using non-linguistic stimuli. The results provide partial support for the temporal sampling framework (Goswami, 2011) in that the quality of CT at low frequencies is related to later reading acquisition. In particular, they add novel evidence regarding the differential role played by delta vs. theta tracking in reading acquisition. Our findings suggest that, while prereaders show CT at both delta and theta rates, only CT at delta rate is related to reading acquisition. This results go in line with a developmental perspective of cortical tracking were delta CT relevance is larger during infancy and prereading stages (Attaheri et al., 2020). This could respond to the fact that spectral energy in the delta band is larger in child directed speech than in adult directed speech (Pérez-Navarro et al., 2022).

It should be noted that we did not test the contribution of delta vs. theta in a single model, given the modest sample size and the fact that a different set of electrodes was used for comparing groups at each frequency, based on the data-driven analysis approach. Interpretations should therefore be performed with caution. However, the characteristics of theta CT suggest that the lack of a role for theta in the CT-reading link is a robust finding. Theta CT was both stronger and much more widely distributed than delta CT, suggesting that if there was an effect of theta CT on reading, we would have been able to detect it. Thus, it seems that CT at theta rate, although present in prereaders, is not particularly relevant for reading acquisition. Additionally, available evidence shows that theta CT mainly reflects acoustic (vs. linguistic) processing of the input. For example, Boucher et al. (2009) showed equivalent theta CT in processing tones, nonsense syllables or utterances, and Molinaro and Lizarazu (2017) also found equivalent theta CT in speech processing vs. rotated speech or amplitude modulated noise. These and additional studies underscore theta’s role in general acoustic/perceptual processing (Etard & Reichenbach, 2019; Prinsloo & Lalor, 2020).

With respect to the role that delta CT plays in reading acquisition, the topographical distribution of the observed effect warrants discussion. Of course, spatial interpretations based on sensor-level analysis of the EEG signal should be performed with caution. Our observation of this effect at frontal sites seems more in line with top-down modulations from frontal (including inferior frontal gyrus and precentral gyri) to temporal auditory regions than from bottom-up effect arising in the auditory cortex. This would suggest that the association between CT during kindergarten and reading skills at the end of first grade reported here stems from the quality of the top-down linguistic or attentional modulation of auditory CT, not through delta CT at the auditory level (Ding et al., 2015; Park et al., 2015; Wild et al., 2012).

In sum, the present study shows that cortical tracking of auditory stimuli at prereading stages precedes and predicts future reading skills in prereaders, underscoring the role of cortical tracking for reading acquisition even before any reading skills have emerged. In particular, it highlights the role that delta tracking plays during this developmental stage, which further suggests that slow components in the speech signal, such as prosody, may be particularly salient in the processing of speech input in prereading stages.

